# Sprayable in Situ Forming PEG Hydrogels for Intranasal Drug Delivery

**DOI:** 10.64898/2026.09.17.752479

**Authors:** Luke Zhao, Gregg A. Duncan, Taj Yeruva

## Abstract

Intranasal administration provides a noninvasive route for local and systemic drug delivery, yet its clinical effectiveness remains limited by rapid mucociliary clearance and short drug residence times within the nasal cavity. Existing stimulus-responsive nasal hydrogels seek to address this challenge but rely on physiological triggers such as temperature, pH, or ionic strength to induce gelation, resulting in variable performance due to natural differences in the nasal cavity. Here, we present a polyethylene glycol (PEG)-based hydrogel platform that overcomes these limitations through rapid in situ gel formation that is independent of physiological stimuli. The two liquid precursor solutions remain sprayable during administration and rapidly crosslink upon deposition to form a cohesive hydrogel depot. Although this dual-component design presents challenges for compatibility with conventional nasal spray devices, we demonstrate successful delivery using commercially available dual-barrel spray systems. The hydrogel maintained spray performance and regional nasal deposition comparable to conventional aqueous formulations while significantly reducing mucociliary transport to prolong mucosal residence. Importantly, reduced mucociliary transport was achieved without impairing ciliary function or compromising epithelial barrier integrity, addressing key safety considerations for repeated intranasal administration. In addition, the hydrogel altered epithelial uptake and transport of biologics, suggesting its potential to improve the nasal delivery of macromolecular therapeutics that typically exhibit poor bioavailability. The results of these study establish a versatile and clinically translatable approach for enhancing intranasal drug delivery across a broad range of therapeutic applications.

## Introduction

Intranasal drug delivery is an attractive route for the local and systemic administration of therapeutics as it enables direct delivery to the site of action while providing rapid systemic absorption through the highly vascularized nasal mucosa, thereby avoiding hepatic first-pass metabolism.^**1–3**^ Beyond small molecule drugs, the nasal route has emerged as a promising platform for the delivery of biologics and mucosal vaccines owing to its accessibility and the presence of a highly immunologically active mucosa.^**4–6**^ However, the clinical potential of intranasal delivery remains limited by rapid mucociliary clearance, which typically removes formulations from the nasal cavity within 15–20 minutes, limiting drug residence time and reducing the bioavailability, thereby necessitating frequent dosing.^**7–9**^

To prolong nasal residence, a variety of mucoadhesive and in situ gelling formulations have been developed using viscosity enhancing polymers and stimuli-responsive materials, including chitosan, carbomers, alginate, gellan gum, and poloxamers. Conventional mucoadhesive formulations typically extend residence time from minutes to several hours, while thermoresponsive, pH-responsive, ion-sensitive, and more recently chemically crosslinked hydrogel systems have further prolonged retention from several hours to days, resulting in improved local drug exposure and enhanced immune responses following intranasal vaccination.^**10–18**^ Despite these advances, a fundamental engineering challenge remains, the material properties required to achieve prolonged retention and rapid gel formation often increase formulation viscosity or initiate gelation during administration, potentially compromising atomization, spray plume geometry, and regional nasal deposition. Consequently, achieving efficient spray delivery while simultaneously forming a robust intranasal depot remains a significant engineering challenge.^**19–21**^ Moreover, spray performance, an essential determinant of clinical efficacy and regulatory acceptance is often inadequately characterized relative to gelation and drug release in the development of intranasal hydrogel systems.

An ideal intranasal hydrogel should behave as a low-viscosity liquid during atomization to ensure reproducible spray performance and clinically relevant deposition, yet rapidly transition into a stable, mucoadhesive depot after deposition to slow mucociliary clearance without disrupting normal epithelial function. In our previous work, we developed a rapid in situ forming polyethylene glycol (PEG) hydrogel based on covalent crosslinking between complementary thiol- and ortho-pyridyl disulfide-functionalized PEG precursors and demonstrated its ability to provide sustained local delivery of biologics while preserving their activity.^**22**^ Here, we investigate the potential of this platform for intranasal drug delivery. Unlike conventional stimulus-responsive nasal gels that rely on physiological temperature, pH, or ionic strength to initiate gelation, the PEG precursor solutions remain sprayable during administration and rapidly form a cohesive hydrogel depot after deposition. We demonstrate that the hydrogel maintains spray performance and regional nasal deposition comparable to conventional aqueous sprays, slows mucociliary transport while preserving ciliary function and epithelial barrier integrity, and modulates the uptake and transport of biologics across the airway epithelium. Together, these findings establish a versatile intranasal delivery platform that integrates efficient spray delivery, rapid post-spray gelation, prolonged mucosal residence, and epithelial safety, addressing key translational barriers that have limited existing nasal hydrogel technologies.

## Materials and Methods

### Materials

4-Arm PEG-SH (PEG-4SH) 20 kDa was purchased from Laysan Bio. 4-Arm PEG-OPSS (PEG-4OPSS) 20 kDa was purchased from Creative PEGWorks. Dithiothreitol (DTT) was obtained from Sigma-Aldrich. Tetramethylrhodamine labeled BSA (TRITC-BSA) was purchased from Protein Mods and Fluorescein isothiocyanate conjugated IgG from human serum (FITC-IgG) was purchased from Millipore Sigma. Fluorescent polystyrene nanoparticles (FluoSpheres™) were purchased from Thermo Fisher Scientific. The MAD Nasal spray tip was purchased from Teleflex. The Aptar Pharma nasal spray tip was kindly provided by Aptar Pharma, and the Medmix dual-syringe delivery system (M-System) was provided as a sample by Medmix. All reagents used for human airway epithelial air–liquid interface (ALI) culture were purchased from STEMCELL Technologies.

### Hydrogel preparation and gelation kinetics

Hydrogels were prepared as described in our previous work.^**22**^ Briefly, PEG-4SH and PEG-4OPSS were separately dissolved in citrate buffer (pH 5.0–6.0), phosphate buffer (pH 5.0–7.5), or Tris buffer (pH 7.0–7.5). Equal volumes of PEG-4SH and PEG-4OPSS solutions prepared in the same buffer were mixed to obtain final polymer concentrations of 1% (w/v) PEG-4SH/1.5% (w/v) PEG-4OPSS or 2% (w/v) PEG-4SH/2% (w/v) PEG-4OPSS. Gelation time was determined using the tube inversion method at 37 °C. Briefly, 100 μL of each polymer solution was mixed in a 1.5 mL microcentrifuge tube maintained on a heat block at 37 °C. The tube was inverted at 10-s intervals until the hydrogel no longer flowed along the walls of the tube. Gelation time was defined as the time required to reach this no-flow state. All formulations were evaluated in triplicate.

### Spray characterization

Spray pattern analysis was performed to evaluate the effect of hydrogel formation on atomization. Phosphate-buffered saline (PBS, pH 7.0) was used as liquid control, while the hydrogel formulation consisted of 2% w/v PEG-4SH and 2% w/v PEG-4OPSS prepared in pH 6.5 phosphate buffer close to average nasal pH of 6.3. Rhodamine B (0.02 mg/mL) was added to all formulations to facilitate visualization of the spray pattern. For hydrogel formulations, PEG-4SH and PEG-4OPSS solutions were loaded separately into two 1 mL syringes (200 μL per syringe) mounted in a Twin-Syringe Delivery System 1:1 (M-System; medmix, Switzerland) fitted with a MAD Nasal mucosal atomization device (Teleflex, NC, USA). The syringe assembly was secured in a vertical orientation, and filter paper was positioned perpendicular to the spray nozzle at a distance of 7 cm as shown in **Figure S1**. Formulations were manually actuated to generate spray patterns on the filter paper. PBS was sprayed using the same device configuration. Following air drying, spray patterns were imaged (n=5) and analyzed using Fiji (ImageJ, National Institutes of Health, MD, USA). The spray region was identified by color thresholding based on the Rhodamine B signal. The maximum Feret diameter (MaxFeret) and minimum Feret diameter (MinFeret) were measured, and spray ovality was calculated as: Ovality = MaxFeret / MinFeret.

### Three-dimensional nasal cast model

Intranasal deposition studies were performed using a transparent 3D - printed nasal cast replica derived from computed tomography (CT) imaging of a 33-year-old male, originally developed by Warken et al. for patient-specific nasal drug delivery studies.^**23**^ The stereolithography (STL) files were generously provided by the authors, and the nasal cast was fabricated at Terrapin Works (University of Maryland, College Park, MD, USA). The cast consisted of five anatomically separable regions corresponding to the anterior nasal cavity (AN), upper turbinate (UT), middle turbinate (MT), lower turbinate (LT), and nasopharynx (NP), enabling quantification of regional deposition as shown in **Figure 2D**.

**Figure 1.**
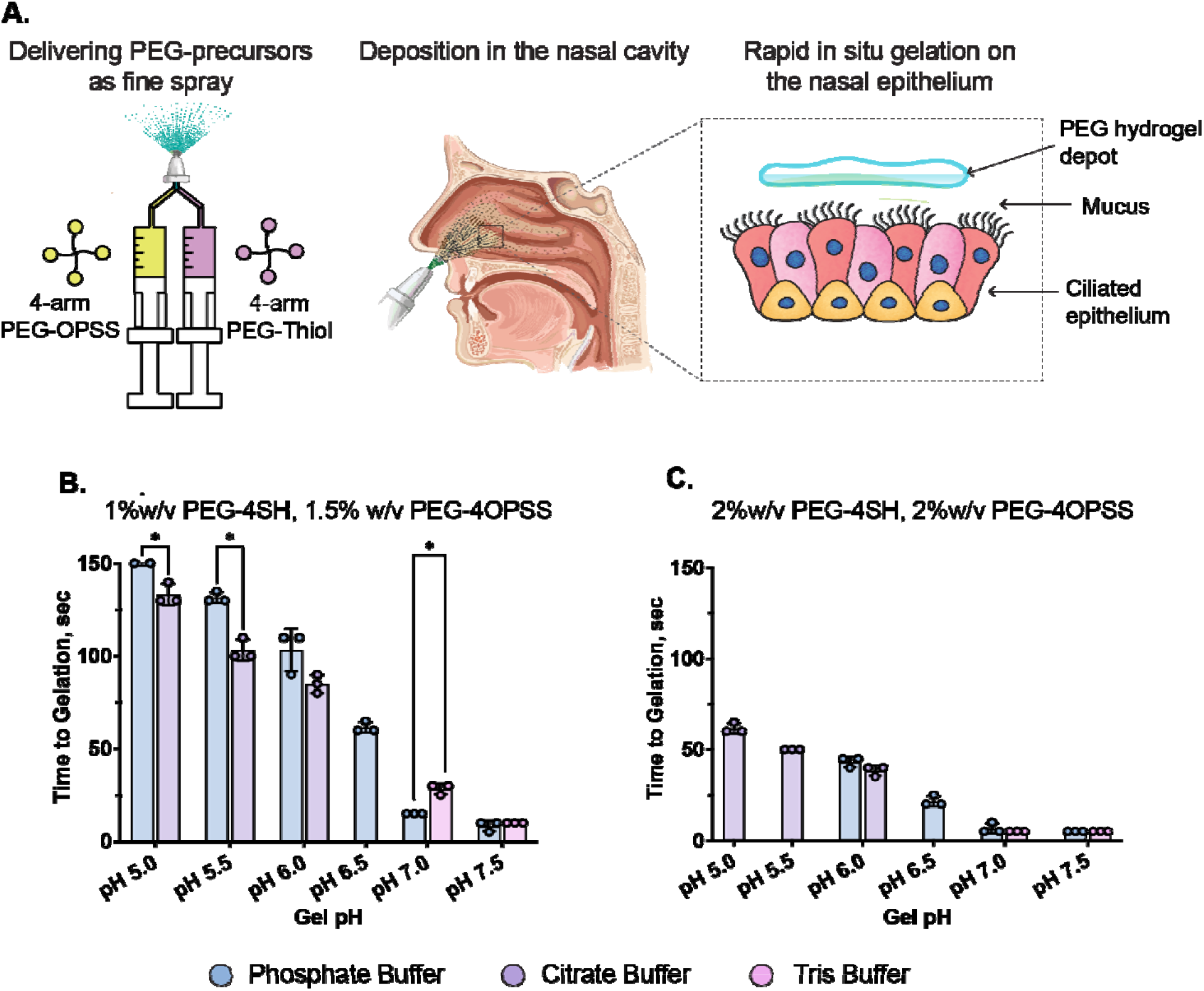
Rapid in situ forming PEG hydrogels for intranasal delivery. (A) Conceptual overview of sprayable rapid in situ forming PEG hydrogels for intranasal delivery. Gelation kinetics of PEG hydrogel formulated with (B) 1% PEG-4SH and 1.5% PEG-4OPSS and (C) 2% PEG-4SH and 2% PEG-4OPSS under varying pH and buffer conditions at 37°C. n=3, *p<0.05 as per unpaired Welch’s t-tests with Holm– Šídák correction for multiple comparisons. Data are presented as mean ± SD.

**Figure 2.**
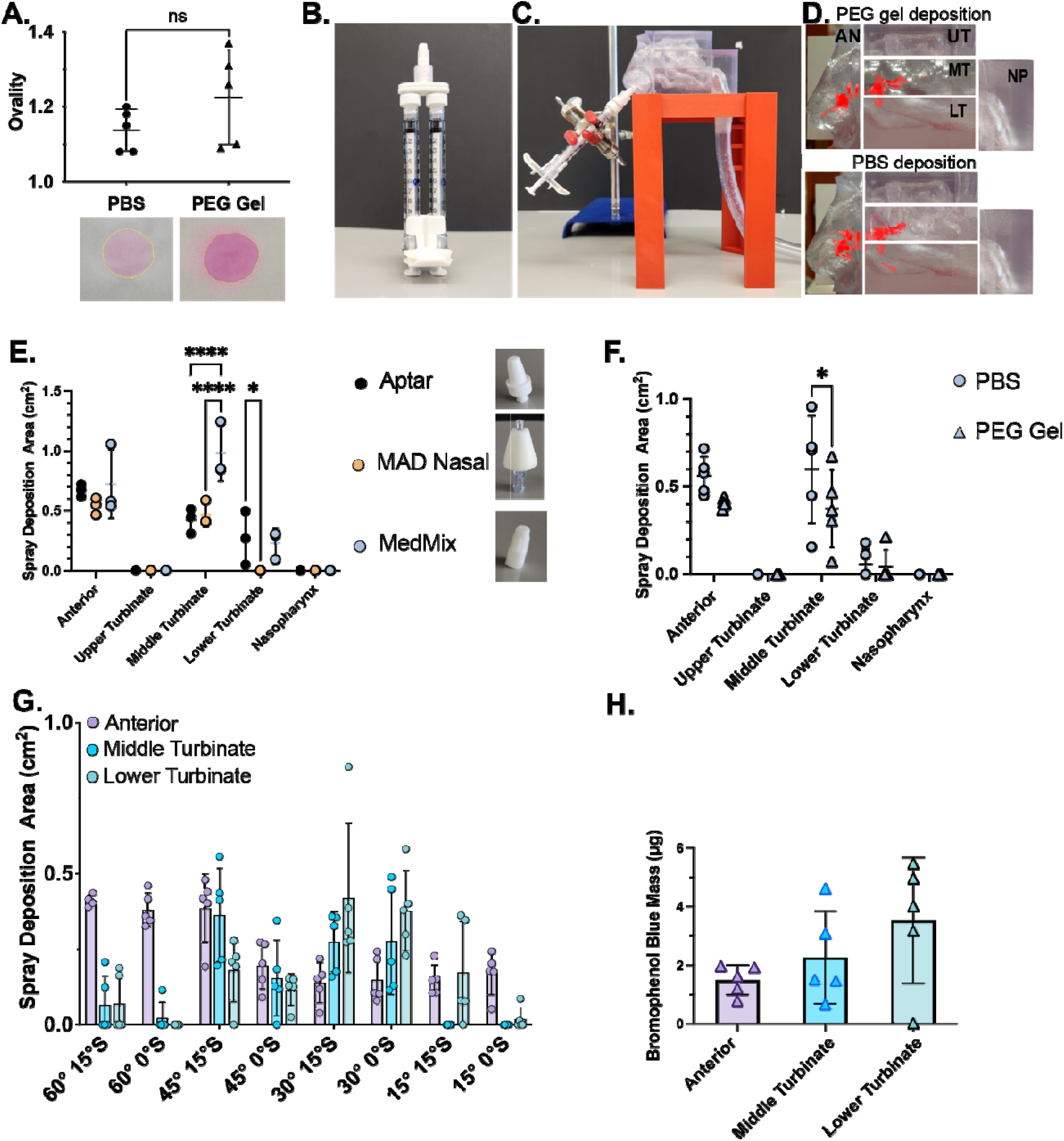
Spray characterization and nasal deposition of rapid in situ forming PEG hydrogels. (A) Quantification of spray ovality and representative spray patterns of PBS and PEG hydrogel formulations deposited onto filter paper following actuation through a twin-syringe delivery system. n=5, nonsignificant (ns) as per unpaired two-tailed Welch’s t-test. (B) Image of the twin-syringe delivery system used for simultaneous administration of PEG-4SH and PEG-4OPSS precursor solutions. (C) Experimental setup for nasal deposition studies using a transparent 3D-printed anatomically accurate nasal cast model. (D) Representative deposition images following administration of PBS or PEG formulations into the nasal cast. Red overlays indicate regions identified during image threshold analysis. Nasal regions analyzed included anterior region (AN), upper turbinate (UT), middle turbinate (MT), lower turbinate (LT), and nasopharynx (NP). (E) Comparison of regional deposition patterns obtained using different commercial spray tip configurations. (F) Comparison of regional nasal deposition profiles between PBS and PEG hydrogel formulations following administration at a 45° coronal angle with 15° sagittal orientation. (G) Regional deposition following PBS administration at varying delivery angles. (H) Mass quantification of deposited PEG hydrogel within the nasal cast following administration at a 30° coronal angle with 15° sagittal orientation. *p<0.05 and ****p<0.0001for two-way ANOVA with Šídák’s multiple comparisons test. Data are presented as mean ± SD.

### Nasal cast experimental setup

Prior to each experiment, the nasal cast was assembled and sealed with vacuum grease and transparent packing tape to prevent air leakage. The nasopharyngeal outlet was connected to a vacuum source operating at a constant flow rate of 20 L min□^**1**^ to simulate inspiratory airflow as shown in **Figure 2C**. Proper sealing was verified by occluding the nostrils and confirming negative pressure within the cast. A Twin-Syringe Delivery System 1:1 (M-System; medmix, Haag, Switzerland) fitted with an interchangeable atomization spray tip was mounted on a laboratory stand as shown in **Figure 2A and 2B** to provide reproducible positioning of the spray device. The stand allowed adjustment of the administration angle relative to both the coronal plane (15°, 30°, 45°, or 60°) and the sagittal plane (0° or 15°). All sprays were manually actuated, and the vacuum was maintained for an additional 30 s following spray administration to allow complete aerosol transport through the nasal cast. Following spray administration, the assembled nasal cast was imaged using a smartphone camera (Galaxy S22, Samsung Electronics). The camera position and imaging distance were maintained constant for all experiments to ensure consistent image acquisition. Images were subsequently cropped to the boundaries of the nasal cast and analyzed in Fiji. Prior to quantitative analysis, images were calibrated to the known dimensions of the nasal cast (11 cm × 6 cm).

### Selection of the atomization spray tip

Three commercially available nasal spray tips (Aptar, MAD Nasal, and medmix) were compared to identify the device providing the most favorable regional deposition profile. PBS (pH 7.0) containing green food coloring was used as the model formulation. Approximately 0.5 mL of formulation was loaded into each device and primed prior to use to eliminate variability associated with connector and spray-tip dead volume. Sprays were administered at an insertion angle of 45° relative to the coronal plane and 15° relative to the sagittal plane. Regional deposition was quantified from digital images using Fiji. Spray deposition was segmented using automated color thresholding, and the deposition area within each anatomical region was quantified (n = 3).

### Comparison of PBS and PEG gel intranasal deposition

Following selection of the medmix Twin-Syringe Delivery System, intranasal deposition of PBS (pH 7.0) and the PEG hydrogel precursor formulation consisting of 2% w/v PEG-4SH and 2% w/v PEG-4OPSS prepared in pH 6.5 phosphate buffer was compared. Green food coloring (6 drops/10 mL) was incorporated into each formulation to facilitate visualization of regional deposition. To account for the device dead volume (∼140 μL, including the connector and spray tip), each syringe was loaded with 120 μL of solution, resulting in a final delivered spray volume of 100 μL. Sprays were administered at an insertion angle of 45° relative to the coronal plane and 15° relative to the sagittal plane. Five independent sprays were performed for each formulation (n = 5). Following spray administration, regional deposition patterns were imaged and quantified using the Fiji (ImageJ) image analysis method described above. Between experiments, the nasal cast was cleaned to remove residual formulation. Following PBS deposition studies, the cast was rinsed thoroughly with tap water. Following hydrogel deposition studies, the cast components were immersed in 40 mM dithiothreitol (DTT) solution to dissolve the disulfide-crosslinked hydrogel, rinsed with water, and allowed to air dry before subsequent use.

### Optimization of administration angle

Following selection of the medmix Twin-Syringe Delivery System, the effect of administration angle on intranasal deposition was evaluated using PBS containing green food coloring. Sprays were administered at coronal angles of 15°, 30°, 45°, and 60°, each evaluated at sagittal orientations of 0° and 15°. Regional deposition was quantified using image-based analysis of the five anatomical regions of the nasal cast. The administration angle providing the maximized deposition within the turbinate regions with minimal anterior deposition was selected for subsequent studies.

### Mass based regional deposition

Regional deposition of the PEG hydrogel formulation 2% w/v PEG-4SH and 2% w/v PEG-4OPSS prepared in pH 6.5 phosphate buffer was quantified at the optimized administration angle (30° relative to the coronal plane and 15° relative to the sagittal plane). Bromophenol blue was incorporated into the hydrogel precursor solutions as a tracer dye. Following spray administration, the nasal cast was disassembled into its five anatomical regions, and deposited formulations were recovered by dissolution in DTT. Bromophenol blue concentration was quantified by UV–visible spectrophotometry using a calibration curve.

### Air–Liquid Interface (ALI) culture of human airway epithelial cells

Air–liquid interface (ALI) cultures were established as previously described.^**24,25**^ Briefly, the hTERT-immortalized human airway epithelial basal cell line BCi-NS1.1 (provided by Dr. Ronald Crystal, Weill Cornell Medical College, New York, NY, USA) was expanded in T-75 tissue culture flasks using PneumaCult™-Ex Plus medium at 37 °C in a humidified atmosphere containing 5% CO□. Cells were seeded (1 × 10□ cells per insert) onto rat tail Type I collagen (50 μg/mL)-coated 6.5 mm Transwell® inserts (0.4 μm pore size; Corning Inc., Corning, NY, USA). Upon reaching confluency, the apical medium was removed, and the basolateral medium was replaced with PneumaCult™-ALI medium to establish ALI cultures. Cells were differentiated for at least 28 days with basolateral medium replaced every other day, resulting in a pseudostratified, mucus-producing, ciliated airway epithelium. Accumulated mucus was removed from the apical surface twice weekly by incubating with 250 μL Dulbecco’s phosphatebuffered saline (DPBS) without calcium and magnesium for 30 min at 37 °C before aspiration.

### Hydrogel application to human airway epithelial cultures

Unless otherwise specified, hydrogel treatments were performed by sequential application of the two precursor solutions onto the apical surface of differentiated HAE cultures. Twenty microliters of the 4% w/v PEG-4SH solution prepared in PBS (pH 7.4) was first applied to the center of the epithelial surface, followed immediately by 20 μL of the 4% w/v PEG-4OPSS solution. prepared in PBS (pH 7.4). Transwell inserts were then gently tilted in multiple directions to promote uniform mixing of the precursor solutions and in situ hydrogel formation consisting of 2% (w/v) PEG-4SH and 2% (w/v) PEG-4OPSS across the epithelial surface.

### Epithelial barrier integrity

The integrity of the airway epithelial barrier following PEG hydrogel treatment was assessed by measuring transepithelial electrical resistance (TEER) and ZO-1 immunofluorescence. TEER was measured using a Millicell® ERS-2 Electrical Resistance System (MilliporeSigma, USA). For baseline measurements, fully differentiated ALI cultures were equilibrated at room temperature for 15 min with 500 μL Dulbecco’s phosphate-buffered saline (DPBS) in the apical compartment and 1 mL DPBS in the basolateral compartment. Following measurement, DPBS was removed, fresh differentiation medium was added to the basolateral compartment, and cultures were returned to the incubator. 24 hours later, cultures were treated with 40 μL of PEG hydrogel precursor formulation. TEER measurements were repeated at 30 min and 24 h post treatment using the same procedure. Three resistance measurements were recorded for each transwell insert, and the average value was used for analysis. The same ALI cultures were used for baseline and post treatment measurements, allowing changes in TEER to be assessed within the same biological replicate over time. TEER values were corrected by subtracting the blank resistance and multiplying by the membrane surface area (0.33 cm^**2**^) and are reported as Ω·cm^**2**^.

Following the final TEER measurement, differentiated ALI cultures were fixed with 4% paraformaldehyde for 15 min at room temperature, washed with PBS, and blocked with 3% bovine serum albumin (BSA) in PBS for 1 h. Cultures were incubated overnight at 4 °C with Alexa Fluor™ 488-conjugated anti-ZO-1 monoclonal antibody (ZO1-1A12, Invitrogen; 1:2000 dilution) prepared in 1% BSA in PBS. After washing with PBS, transwell membranes were excised, mounted onto glass microscope slides, and imaged using a Zeiss LSM 800 confocal microscope

### Mucociliary transport and ciliary beat frequency

Mucociliary transport (MCT) and ciliary beat frequency (CBF) were quantified as previously described.^**24**^ Briefly, 2 μm red fluorescent microspheres were sonicated for 10 min, diluted 1:2000 in sterile phosphate-buffered saline (PBS), and mixed thoroughly. 1 μL of the microsphere suspension was added to 100 μL of either PBS or the PEG hydrogel precursor formulation. To establish baseline MCT, accumulated mucus was removed from differentiated ALI cultures by washing the apical surface with DPBS. 24 hours later, 40 μL of PBS containing fluorescent microspheres was applied to the apical surface, and baseline MCT was recorded by fluorescence video microscopy. Following imaging, the apical surface was washed with DPBS, and the cultures were returned to the incubator. 24 hours later, 40 μL of the PEG hydrogel precursor formulation containing fluorescent microspheres was applied to the same cultures, and MCT was measured at 30 min and 24 h post-treatment. Videos were acquired from five random fields of view per transwell insert using a 10× objective. Each field was recorded for 20 s with an exposure time of 150 ms. Microsphere trajectories were analyzed using a custom MATLAB particle-tracking algorithm, and the median particle velocity from each video was used for statistical analysis.

Immediately following MCT imaging, the microscope was switched to brightfield mode, and the focus on the epithelium was adjusted until ciliary motion was clearly visible. Brightfield videos were acquired from three randomly selected fields of view per transwell insert using 10× objective for 10 s with an exposure time of 20 ms. CBF was quantified using Fiji and a custom MATLAB analysis script. Briefly, mean pixel intensity over time was extracted from three randomly selected regions of interest within each video, and the number of local maxima over the time corresponding to ciliary beating were identified to calculate CBF. The mean CBF value from each video was plotted.

### BSA transport across HAE cultures

Transport of a model protein across differentiated human airway epithelial (HAE) cultures was evaluated using tetramethylrhodamine isothiocyanate-labeled bovine serum albumin (TRITCBSA). TRITC-BSA was prepared at a final concentration of 0.1% (w/v) in phosphate-buffered saline (PBS) or incorporated into the PEG hydrogel. Forty microliters of each formulation were applied to the apical surface of differentiated HAE cultures as described above. At the predetermined time points, basolateral medium was collected and replaced with fresh PneumaCult-ALI medium. TRITC-BSA transport was quantified using a fluorescence plate reader (excitation: 545 nm; emission: 575 nm), and concentrations were determined from a standard curve prepared in PneumaCult-ALI medium.

### IgG transport, cellular uptake, and barrier integrity

Differentiated HAE cultures were treated with PBS, FITC-IgG in PBS, or FITC-IgG incorporated into the PEG hydrogel. Forty microliters of each formulation were applied to the apical surface of the cultures as described above. Epithelial barrier integrity was assessed by measuring transepithelial electrical resistance (TEER) before treatment and at 24 h post treatment as described above. FITC-IgG transport across the epithelial barrier was quantified using the procedure described for BSA transport (excitation: 490 nm; emission: 525 nm). To quantify cellular uptake, cultures were washed with PBS to remove unbound FITC-IgG, fixed with 4% paraformaldehyde, counterstained with DAPI, and mounted onto glass microscope slides. Fluorescence images were acquired using a Zeiss LSM 800 confocal microscope under identical acquisition settings for all samples. Five randomly selected fields of view were acquired per transwell insert, and intracellular FITC-IgG uptake was quantified in Fiji by measuring the mean fluorescence intensity. DAPI staining was used for visualization only and was not included in the quantitative analysis.

## Results and Discussion

### Designing rapid in situ forming PEG hydrogels for intranasal administration

**Figure 1A** illustrates the design strategy for the rapid in situ forming PEG hydrogel platform for intranasal drug delivery. Two precursor solutions consisting of thiol-terminated four-arm polyethylene glycol (PEG-4SH) and ortho-pyridyl disulfide-terminated four-arm polyethylene glycol (PEG-4OPSS) are loaded into a twin-syringe delivery device at a 1:1 ratio and mixed immediately before atomization. Following intranasal administration, the precursor solutions deposit throughout the nasal cavity and crosslink in situ to form a mucoadhesive hydrogel depot on the airway epithelium, enabling prolonged local retention and sustained therapeutic delivery.

In our previous work, we developed this PEG hydrogel platform and demonstrated gelation in < 30 s under physiological conditions (37 °C, pH 7.4 in phosphate buffer).^**22**^ Here, we evaluated gelation behavior under conditions relevant for intranasal drug delivery. The nasal cavity is mildly acidic (∼ pH 5.5 - 6.5) while inflammatory conditions such as allergic rhinitis can increase nasal pH to approximately 7.2–8.3.^**26,27**^ In addition, commercially available intranasal products span a broad range of formulation pH values and buffering systems.^**28**^ Therefore, gelation kinetics of two PEG hydrogel compositions (1% w/v PEG-4SH/1.5% w/v PEG-4OPSS and 2% w/v PEG-4SH/2% w/v PEG-4OPSS) were evaluated in citrate (pH 5.0–6.0), phosphate (pH 5.0– 7.5), and Tris (pH 7.0–7.5) buffers. Both formulations exhibited pH-dependent gelation kinetics, with gelation time decreasing as pH increased while remaining largely independent of buffer composition (**Fig. 1B, C**). As expected, the lower concentration formulation consistently exhibited longer gelation times than the 2% w/v PEG-4SH/2% w/v PEG-4OPSS formulation.

The 2% w/v PEG-4SH/2% w/v PEG-4OPSS formulation prepared in phosphate buffer at pH 6.5 gelled within approximately 25 s, providing sufficient working time for spray administration while enabling rapid in situ hydrogel formation following deposition under physiologically relevant nasal conditions. This formulation was therefore selected for all subsequent studies.

### Rapid forming PEG gels preserve sprayability and nasal deposition

Efficient intranasal drug delivery requires formulations to generate reproducible spray patterns that promote deposition within the turbinate region of the nasal cavity, where most drug absorption occurs. Because spray pattern is a critical quality attribute influencing dose uniformity and regional deposition, spray performance was quantified using spray ovality. Ovality was defined as the ratio of the maximum to minimum spray diameters, with a value of 1 indicating a perfectly circular spray pattern and increasing values reflecting greater spray asymmetry.^**29,30**^ The PEG hydrogel formulation produced spray patterns comparable to the PBS control, with no significant differences in spray ovality between formulations (**Fig. 2A**). Both formulations generated near-circular spray patterns (ovality ≈1), demonstrating that incorporation of the PEG precursors did not adversely affect spray atomization or symmetry compared to liquid formulations.

To determine whether the PEG hydrogel maintained deposition characteristics following atomization, regional deposition was evaluated using a twin-syringe delivery system in a 3D printed anatomically accurate nasal cast model (**Fig. 2B–D**). Consistent with previous reports, negligible deposition was observed in the upper turbinate and nasopharyngeal regions under all tested conditions.^**20,31**^ Among commercial spray tips evaluated, Medmix and Aptar tips produced comparable deposition profiles (**Fig. 2E**). The Medmix tip was selected for subsequent studies because of its compact geometry.

Regional deposition of the PEG hydrogel was then compared with PBS using the Medmix spray tip at a 45° coronal (C) and 15° sagittal (S) administration angle. Both formulations exhibited similar deposition profiles throughout the nasal cavity, indicating that hydrogel formation did not compromise spray distribution following administration (**Fig. 2F**). To further optimize intranasal delivery, multiple administration angles were evaluated using PBS. Administration at 30°C, 0°S and 30°C, 15°S produced greater deposition within the lower and middle turbinates while reducing deposition in the anterior region compared with other orientations tested (**Fig. 2G**). Incorporation of the mixing chamber did not alter nasal deposition at the optimized 30° C, 15° S orientation (**Fig. S2**), supporting its compatibility with intranasal delivery. Absorbance based mass quantification of PEG gel showed deposition trends consistent with imaging analysis, demonstrating the highest deposition in the lower turbinates, followed by the middle turbinates and anterior region (**Fig. 2H**). Collectively, these findings demonstrate that the rapid in situ forming PEG hydrogel retains the spray atomization and regional deposition characteristics of conventional liquid nasal sprays despite undergoing rapid crosslinking after administration.

### Rapid forming PEG gel slows mucociliary clearance without compromising epithelial barrier function

Mucociliary clearance (MCC) is a major barrier to effective intranasal drug delivery, with deposited materials typically cleared from the nasal cavity within ∼15 to 20 min. While this defense mechanism protects the respiratory tract from inhaled pathogens and particulates, it also limits the absorption window for locally administered therapeutics. Therefore, an ideal intranasal depot should slow mucociliary transport without disrupting normal epithelial function. To evaluate epithelial compatibility, differentiated BCi-NS1.1 human airway epithelial (HAE) cultures grown at an air-liquid interface (ALI) were used, as they recapitulate the pseudostratified airway epithelium, including functional ciliated cells, mucus production, and coordinated mucociliary transport. Compared with untreated controls, PEG hydrogel treatment significantly reduced mucociliary transport 24 hr. after administration (**Fig. 3B & 3C**), consistent with the formation of a surface-associated hydrogel depot that resists rapid clearance. Importantly, this reduction in particle transport was not accompanied by impairment of ciliary function, as ciliary beat frequency remained unchanged 24 hr. following treatment (**Fig. 3D**). Likewise, epithelial barrier integrity was preserved, with no significant changes in transepithelial electrical resistance (TEER) (**Fig. 3E**) or ZO-1 tight junction organization (**Fig. 3F**). Together, these findings demonstrate that the PEG hydrogel slows mucociliary transport while maintaining normal ciliary activity and epithelial barrier integrity, supporting its suitability as a biocompatible intranasal depot for sustained local drug delivery.

**Figure 3.**
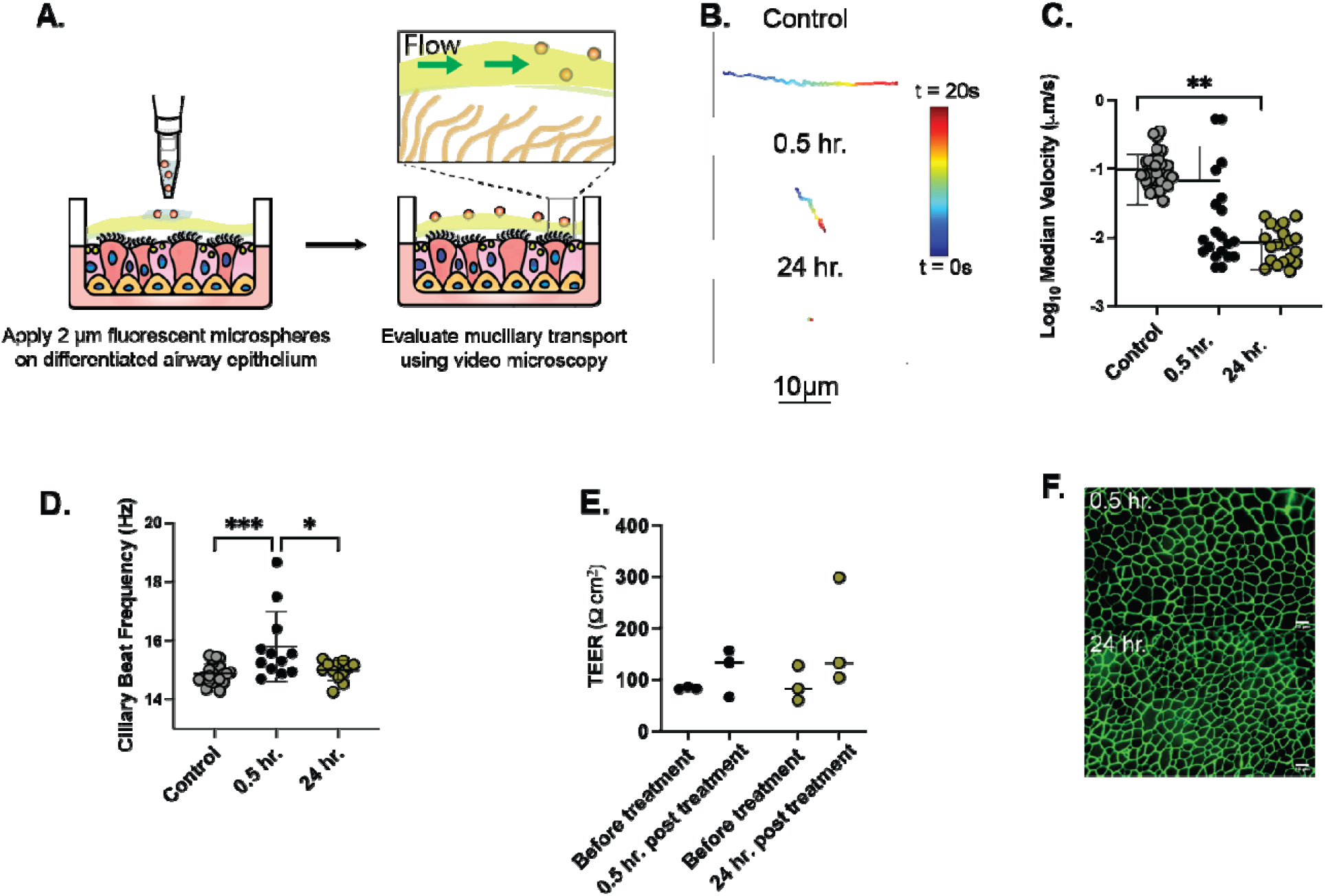
In vitro human airway epithelial compatibility to rapid in situ forming PEG hydrogels. (A) Schematic overview of MCT workflow. *Adapted from Song et al*., *Sci. Adv. 2022*.^*25*^ (B) Representative trajectories of single microsphere over time (C) Log_10_ of median velocities of tracked microspheres from each video (n = 4 cultures per group; n = 5 videos per culture) (B) Ciliary beat frequency (CBF) (n = 4 cultures per group; n = 3 videos per culture) (C) transepithelial electrical resistance (TEER) (n = 3 cultures per group) *p<0.05, ***p<0.001 for one-way ANOVA with Tukey’s multiple comparison test, and (D) representative ZO-1 immunofluorescence images of differentiated BCi-NS1.1 airway epithelial ALI cultures at 0.5 hr. and 24 hr. following treatment with PEG hydrogels. Scale bars = 10 μm.

### Rapid forming PEG gels facilitate controlled release of biologics and enhanced uptake at airway epithelium

To evaluate the potential of the PEG hydrogel platform for intranasal delivery of biologics, the transepithelial transport of TRITC-BSA (66 kDa) and FITC-IgG (150 kDa) was evaluated using differentiated BCi-NS1.1 human airway epithelial (HAE) air-liquid interface (ALI) cultures. Compared with PBS, delivery from the PEG hydrogel significantly increased BSA transport across the epithelial layer from ∼ 30% to 80% over 24 hr. (**Fig. 4A**). Albumin is internalized through endocytosis and can subsequently engage FcRn-mediated intracellular trafficking, which protects albumin from degradation and facilitates its recycling or transcytosis.^**32**^ The prolonged epithelial residence provided by the hydrogel likely increased opportunities for epithelial uptake and transport of BSA.

**Figure 4.**
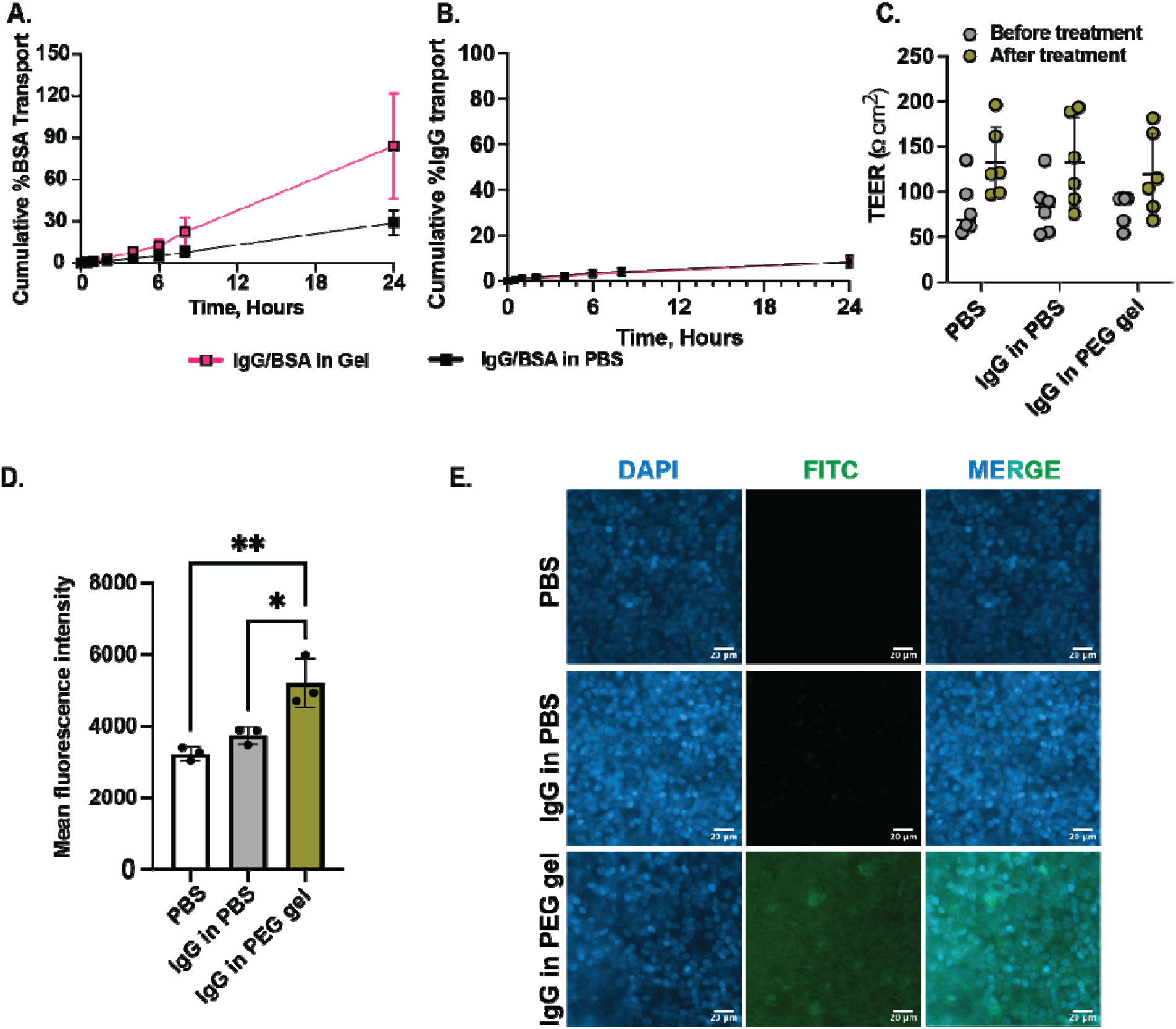
Transport and epithelial uptake of biologics delivered from rapid in situ forming PEG hydrogels across airway epithelial air-liquid interface (ALI) cultures. Quantification of (A) TRITCBSA transport (n=6 cultures per group), (B) FITC-IgG transport (n=6 cultures per group), (C) Tran epithelial electrical resistance (TEER) (n=6 cultures per group), (D) Intracellular FITC-IgG uptake quantified as mean fluorescence intensity (n=3 cultures per group), and (E) Representative fluorescence images showing FITC-IgG uptake by differentiated airway epithelial cultures following apical administration in PBS or PEG hydrogel formulations. *p<0.05 and **p<0.01 for one-way ANOVA with Tukey’s multiple comparison test. Differences in TEER values are non-significant before and after treatment for two-way ANOVA with with Šídák’s multiple comparisons test. Data are presented as mean ± SD.

In contrast, transepithelial transport of FITC-IgG remained below 10% irrespective of the delivery vehicle (**Fig. 4B**), despite the PEG hydrogel significantly increasing epithelial-associated IgG fluorescence compared with PBS controls (**Fig. 4D, E**). Because airway epithelial cells express FcRn, increased epithelial association may reflect enhanced cellular uptake resulting from prolonged apical exposure and subsequent intracellular trafficking.^**33–35**^ However, the lack of increased basolateral transport suggests that internalized IgG was predominantly retained within epithelial cells or recycled rather than undergoing efficient transcytosis. This observation is consistent with previous studies demonstrating that FcRn-mediated IgG transport across the airway epithelium is relatively limited despite functional receptor expression. Importantly, epithelial barrier integrity remained intact following treatment, as evidenced by unchanged TEER values across all groups (Fig. 4C), indicating that the increased epithelial association of IgG was not attributable to compromised epithelial junctions. Collectively, these findings demonstrate that rapid in situ forming PEG hydrogels differentially regulate the epithelial fate of biologics in a cargo-dependent manner. By enhancing albumin transport while promoting epithelial association of IgG without compromising barrier integrity, this platform provides prolonged mucosal exposure to therapeutic proteins while preserving normal epithelial function.

## Conclusions

This work presents a new design for intranasal drug delivery by decoupling sprayability from gel formation. The rapid in situ forming PEG hydrogel enables efficient atomization and clinically relevant nasal deposition before rapidly forming a stable mucoadhesive depot that prolongs mucosal residence without compromising ciliary function and epithelial barrier integrity. By overcoming key translational barriers associated with conventional nasal gels, this platform provides a broadly adaptable strategy for the sustained intranasal delivery of small molecules, biologics, and vaccines.

## Acknowledgements

This study was supported by the UM Ventures Medical Device Development Fund grant to GAD.

## References

(1) Xu, D.; Song, X.-J.; Chen, X.; Wang, J.-W.; Cui, Y.-L. Advances and Future Perspectives of Intranasal Drug Delivery: A Scientometric Review. Journal of Controlled Release 2024, 367, 366–384. 10.1016/j.jconrel.2024.01.053

(2) Costantino, H. R.; Illum, L.; Brandt, G.; Johnson, P. H.; Quay, S. C. Intranasal Delivery: Physicochemical and Therapeutic Aspects. International Journal of Pharmaceutics 2007, 337 (1), 1–24. 10.1016/j.ijpharm.2007.03.025

(3) Keller, L.-A.; Merkel, O.; Popp, A. Intranasal Drug Delivery: Opportunities and Toxicologic Challenges during Drug Development. Drug Delivery and Translational Research 2022, 12 (4), 735–757. 10.1007/s13346-020-00891-5

(4) Kiyono, H.; Ernst, P. B. Nasal Vaccines for Respiratory Infections. Nature 2025, 641 (8062), 321–330. 10.1038/s41586-025-08910-6

(5) Ramvikas, M.; Arumugam, M.; Chakrabarti, S. R.; Jaganathan, K. S. Chapter Fifteen - Nasal Vaccine Delivery. In Micro and Nanotechnology in Vaccine Development; Skwarczynski, M., Toth, I., Eds.; William Andrew Publishing, 2017; pp 279–301. 10.1016/B978-0-323-39981-4.00015-4

(6) Pagni, R. L.; Cunha-Neto, E.; Silva Santos, Y. da; Postól, E.; Alencar, R. E. de; Moretti, A.; Santos, J. J.; Silva, T. L.; Daher, I. P.; Knirsch, M. C.; Nunes, J. P. S.; Toma, S. H.; Araki, K.; Remuzgo, C.; Jacintho, L. C.; Demarchi, L. M.; de Oliveira, V. L.; Coelho, V.; Boscardin, S. B.; Rosa, D. S.; Batalha-Carvalho, J. V.; Moro, A. M.; Santos, K. S.; Stephano, M. A.; Kalil, J.; Ares, A. C.; Takara, A. C. K. K.; de Oliveira, J. R.; Yamamoto, M. M.; Marin, M. L. C.; Benedetti, P. R.; Almeida, R. R.; de Barros, S. F.; Monteiro, S. M.; Palacios, S. A.; dos Santos, S. R.; da Silva, W. R.; On behalf of COVID-19 Brazil Team. An Innovative Nasal Nanovaccine against SARS-CoV-2 Induces Systemic and Upper Airway Immunity Controlling Viral Replication. npj Vaccines 2026, 11 (1), 82. 10.1038/s41541-026-01407-x

(7) Agu, R. U. Challenges in Nasal Drug Absorption: How Far Have We Come? Therapeutic Delivery 2016, 7 (7), 495–510. 10.4155/tde-2016-0022

(8) Shah, S. A.; Berger, R. L.; McDermott, J.; Gupta, P.; Monteith, D.; Connor, A.; Lin, W. Regional Deposition of Mometasone Furoate Nasal Spray Suspension in Humans. Allergy and Asthma Proceedings 2015, 36 (1), 48–57. 10.2500/aap.2015.36.3817

(9) Rogers, T. D.; Button, B.; Kelada, S. N. P.; Ostrowski, L. E.; Livraghi-Butrico, A.; Gutay, M. I.; Esther, C. R.; Grubb, B. R. Regional Differences in Mucociliary Clearance in the Upper and Lower Airways. Frontiers in Physiology 2022, 13. 10.3389/fphys.2022.842592

(10) Wu, Y.; Wei, W.; Zhou, M.; Wang, Y.; Wu, J.; Ma, G.; Su, Z. Thermal-Sensitive Hydrogel as Adjuvant-Free Vaccine Delivery System for H5N1 Intranasal Immunization. Biomaterials 2012, 33 (7), 2351–2360. 10.1016/j.biomaterials.2011.11.068

(11) Zhong, Y.; Su, C.; Wu, S.; Miao, C.; Wang, B. Nasal Delivery of an Immunotherapeutic Vaccine in Thermosensitive Hydrogel against Allergic Asthma. International Immunopharmacology 2023, 116, 109718. 10.1016/j.intimp.2023.109718

(12) Bedford, J. G.; Caminschi, I.; Wakim, L. M. Intranasal Delivery of a Chitosan-Hydrogel Vaccine Generates Nasal Tissue Resident Memory CD8+ T Cells That Are Protective against Influenza Virus Infection. Vaccines 2020, 8 (4), 572.

(13) Varma, D. M.; Batty, C. J.; Stiepel, R. T.; Graham-Gurysh, E. G.; Roque, J. A. I.; Pena, E. S.; Hasan Zahid, M. S.; Qiu, K.; Anselmo, A.; Hill, D. B.; Ross, T. M.; Bachelder, E. M.; Ainslie, K. M. Development of an Intranasal Gel for the Delivery of a Broadly Acting Subunit Influenza Vaccine. ACS Biomater. Sci. Eng. 2022, 8 (4), 1573–1582. 10.1021/acsbiomaterials.2c00015

(14) Fu, W.; Guo, M.; Zhou, X.; Wang, Z.; Sun, J.; An, Y.; Guan, T.; Hu, M.; Li, J.; Chen, Z.; Ye, J.; Gao, X.; Gao, G. F.; Dai, L.; Wang, Y.; Chen, C. Injectable Hydrogel Mucosal Vaccine Elicits Protective Immunity against Respiratory Viruses. ACS Nano 2024, 18 (17), 11200–11216. 10.1021/acsnano.4c00155

(15) Yeruva, T.; Yang, S.; Doski, S.; Duncan, G. A. Hydrogels for Mucosal Drug Delivery. ACS Appl. Bio Mater. 2023, 6 (5), 1684–1700. 10.1021/acsabm.3c00050

(16) Pastor, Y.; Ting, I.; Martínez, A. L.; Irache, J. M.; Gamazo, C. Intranasal Delivery System of Bacterial Antigen Using Thermosensitive Hydrogels Based on a Pluronic-Gantrez Conjugate. International Journal of Pharmaceutics 2020, 579, 119154. 10.1016/j.ijpharm.2020.119154

(17) Zhou, S.; Zheng, X.; Chen, J.; Xu, Y.; Du, X.; Wang, C.; Cui, P.; Qiu, L.; Jiang, P.; Ni, X. Liposome Loaded Ion/Temperature Dual Responsive Gellan Gum Hydrogel as Potential Nasal-To-Brain Delivery System. Journal of Biomedical Nanotechnology 2022, 18 (2), 571–580. 10.1166/jbn.2022.3253

(18) Qian, L.; Cook, M. T.; Dreiss, C. A. In Situ Gels for Nasal Delivery: Formulation, Characterization and Applications. Macromolecular Materials and Engineering 2025, 310 (6), 2400356. 10.1002/mame.202400356

(19) Guo, Y.; Laube, B.; Dalby, R. The Effect of Formulation Variables and Breathing Patterns on the Site of Nasal Deposition in an Anatomically Correct Model. Pharmaceutical Research 2005, 22 (11), 1871–1878. 10.1007/s11095-005-7391-9

(20) Pu, Y.; Goodey, A. P.; Fang, X.; Jacob, K. A Comparison of the Deposition Patterns of Different Nasal Spray Formulations Using a Nasal Cast. Aerosol Science and Technology 2014, 48 (9), 930–938. 10.1080/02786826.2014.931566

(21) Kundoor, V.; Dalby, R. N. Effect of Formulation- and Administration-Related Variables on Deposition Pattern of Nasal Spray Pumps Evaluated Using a Nasal Cast. Pharmaceutical Research 2011, 28 (8), 1895–1904. 10.1007/s11095-011-0417-6

(22) Yeruva, T. K.; Morris, I., Robert J.; Kumar, S.; Zhao, L.; Kofinas, P.; Duncan, G. Rapid in Situ Forming PEG Hydrogels for Mucosal Drug Delivery. Biomater. Sci. 2025. 10.1039/D4BM01101E

(23) Warnken, Z. N.; Smyth, H. D. C.; Davis, D. A.; Weitman, S.; Kuhn, J. G.; Williams, R. O., III. Personalized Medicine in Nasal Delivery: The Use of Patient-Specific Administration Parameters To Improve Nasal Drug Targeting Using 3D-Printed Nasal Replica Casts. Molecular Pharmaceutics 2018, 15 (4), 1392–1402. 10.1021/acs.molpharmaceut.7b00702

(24) Corkran, M.; Boboltz, A.; Duncan, G. A.; Scull, M. A. Methods for Discerning the Impact of Mucus on Host Defenses Against Viral Infection. Current Protocols 2025, 5 (9), e70201. 10.1002/cpz1.70201

(25) Song, D.; Iverson, E.; Kaler, L.; Boboltz, A.; Scull, M. A.; Duncan, G. A. MUC5B Mobilizes and MUC5AC Spatially Aligns Mucociliary Transport on Human Airway Epithelium. Sci. Adv. 2022, 8 (47), eabq5049. 10.1126/sciadv.abq5049

(26) Washington, N.; Steele, R. J. C.; Jackson, S. J.; Bush, D.; Mason, J.; Gill, D. A.; Pitt, K.; Rawlins, D. A. Determination of Baseline Human Nasal pH and the Effect of Intranasally Administered Buffers. International Journal of Pharmaceutics 2000, 198 (2), 139–146. 10.1016/S0378-5173(99)00442-1

(27) England, R. J. A.; Homer, J. J.; Knight, L. C.; Ell, S. R. Nasal pH Measurement: A Reliable and Repeatable Parameter. Clinical Otolaryngology & Allied Sciences 1999, 24 (1), 67–68. 10.1046/j.1365-2273.1999.00223.x

(28) Kulkarni, V. S.; Shaw, C. Chapter 6 - Aerosols and Nasal Sprays. In Essential Chemistry for Formulators of Semisolid and Liquid Dosages; Academic Press: Boston, 2016; pp 71–97. 10.1016/B978-0-12-801024-2.00006-6

(29) Research, C. for D. E. and. Nasal Spray and Inhalation Solution, Suspension, and Spray Drug Products--Chemistry, Manufacturing, and Controls Documentation. https://www.fda.gov/regulatory-information/search-fda-guidance-documents/nasal-spray-and-inhalation-solution-suspension-and-spray-drug-products-chemistry-manufacturing-and (accessed 2026-08-06)

(30) Ehrick, J. D.; Shah, S. A.; Shaw, C.; Kulkarni, V. S.; Coowanitwong, I.; De, S.; Suman, J. D. Considerations for the Development of Nasal Dosage Forms. In Sterile Product Development: Formulation, Process, Quality and Regulatory Considerations; Kolhe, P., Shah, M., Rathore, N., Eds.; Springer New York: New York, NY, 2013; pp 99–144. 10.1007/978-1-4614-7978-9_5

(31) Shah, S. A.; Dickens, C. J.; Ward, D. J.; Banaszek, A. A.; George, C.; Horodnik, W. Design of Experiments to Optimize an In Vitro Cast to Predict Human Nasal Drug Deposition. Journal of Aerosol Medicine and Pulmonary Drug Delivery 2014, 27 (1), 21–29. 10.1089/jamp.2012.1011

(32) Anderson, C. L.; Chaudhury, C.; Kim, J.; Bronson, C. L.; Wani, M. A.; Mohanty, S. Perspective –FcRn Transports Albumin: Relevance to Immunology and Medicine. Trends in Immunology 2006, 27 (7), 343–348. 10.1016/j.it.2006.05.004

(33) Togami, K.; Wolf, W.; Olson, L. C.; Card, M.; Shen, L.; Schaefer, A.; Okuda, K.; Zeitlin, L.; Pauly, M.; Whaley, K.; Pickles, R. J.; Lai, S. K. Impact of mAb-FcRn Affinity on IgG Transcytosis across Human Well-Differentiated Airway Epithelium. Frontiers in Immunology 2024, Volume 15-2024. 10.3389/fimmu.2024.1371156

(34) Roopenian, D. C.; Akilesh, S. FcRn: The Neonatal Fc Receptor Comes of Age. Nature Reviews Immunology 2007, 7 (9), 715–725. 10.1038/nri2155

(35) Ledo, A. M.; Dimke, T.; Tschantz, W. R.; Rowlands, D.; Growcott, E. The Role of Airway Mucus and Diseased Pulmonary Epithelium on the Absorption of Inhaled Antibodies. International Journal of Pharmaceutics 2023, 647, 123519. 10.1016/j.ijpharm.2023.123519

