## Supporting Information for "Sprayable in Situ Forming PEG Hydrogels for Intranasal Drug Delivery"

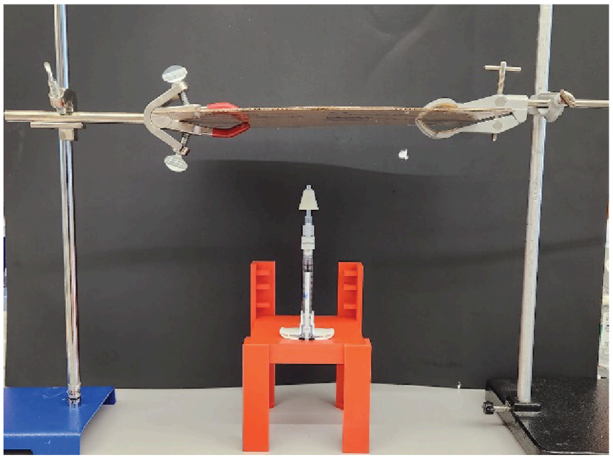

**Figure S1. Experimental setup for spray pattern analysis.** The syringe assembly was secured in a vertical orientation, with filter paper positioned perpendicular to the spray nozzle at 7 cm.

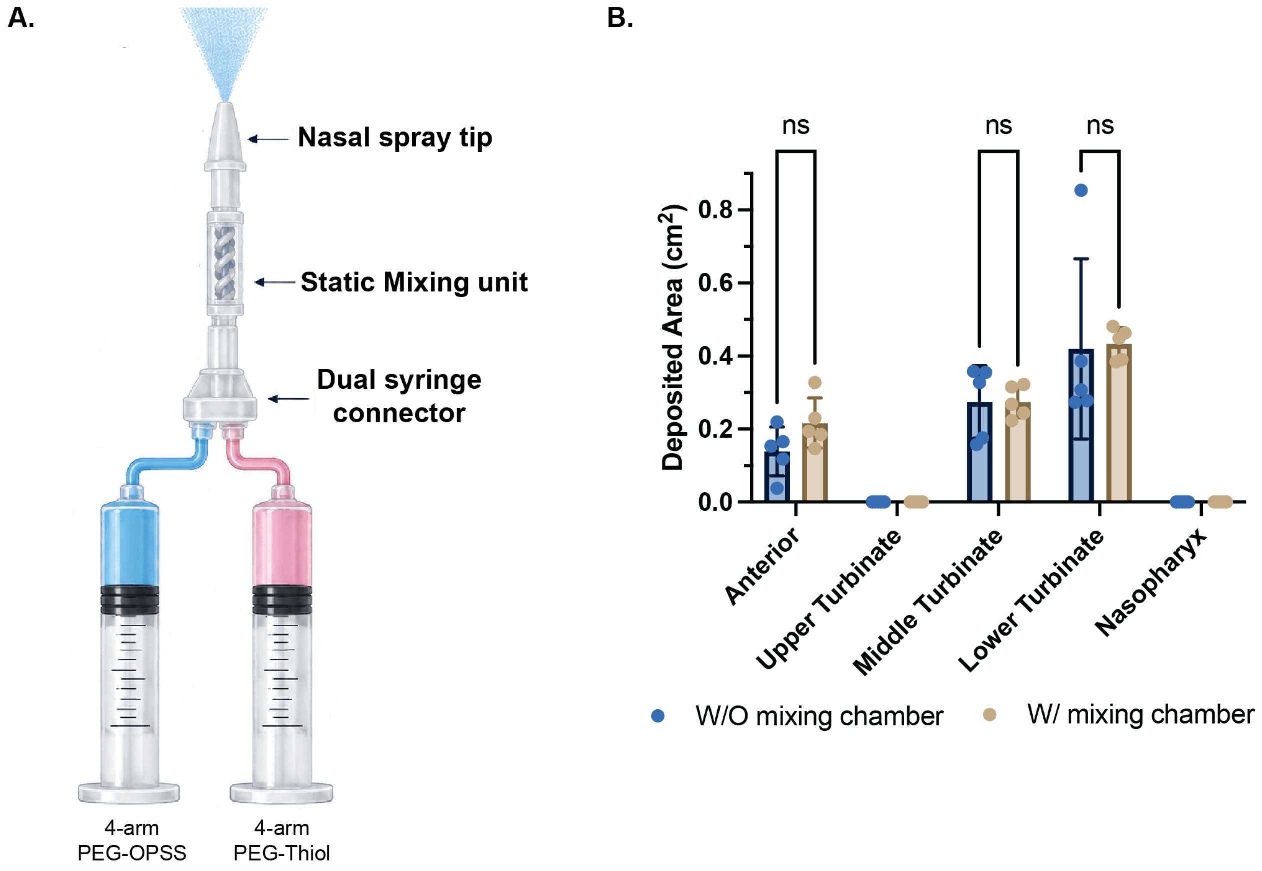


**Figure S2. Effect of the mixing chamber on nasal spray deposition.** (A) Schematic of the medmix twin syringe spray assembly incorporating the connector, mixing chamber, and nasal spray tip. (B) Regional nasal deposition of PBS at the optimized 30° coronal and 15° sagittal administration angle with and without the mixing chamber. Incorporation of the mixing chamber did not significantly alter the regional nasal deposition profile. Data are presented as mean ± SD (n = 5).
